# Massively parallel single-cell profiling of transcriptome and multiple epigenetic proteins in cell fate regulation

**DOI:** 10.1101/2023.04.04.535478

**Authors:** Haiqing Xiong, Qianhao Wang, Chen C. Li, Aibin He

## Abstract

Sculpting the epigenome with a combination of histone modifications and transcription factor (TF) occupancy determines gene transcription and cell fate specification. Here we first develop uCoTarget, utilizing a split-pool barcoding strategy for realizing ultra-high throughput single-cell joint profiling of multiple epigenetic proteins. Through extensive optimization for sensitivity and multimodality resolution, we demonstrate that uCoTarget enables simultaneous detection of five histone modifications (H3K27ac, H3K4me3, H3K4me1, H3K36me3 and H3K27me3) in 19,860 single cells. We applied uCoTarget to the *in vitro* generation of hematopoietic stem/progenitor cells (HSPCs) from human embryonic stem cells, presenting multimodal epigenomic profiles in 26,418 single cells. uCoTarget with high sensitivity per modality reveals establishment of pairing of HSPC enhancers (H3K27ac) and promoters (H3K4me3) along the differentiation trajectory and RUNX1 engagement priming for the H3K27ac activation along the HSPC path. We then develop uCoTargetX, an expansion of uCoTarget to simultaneously measure transcriptome and multiple epigenome targets. Together, our methods enable generalizable, versatile multi-modal profiles for reconstructing comprehensive epigenome and transcriptome landscapes and analyzing the regulatory interplay at single-cell level.

## INTRODUCTION

Cell fate specification and gene expression specificity are largely encoded in the multimodal epigenome, which is characteristic of combinatorial features of various histone modifications and binding of non-histone chromatin proteins, such as chromatin remodelers and transcription factors (TFs)^1–4^. Understanding of the co-existence and regulatory interplay among these molecular modalities is important to predict future gene expression and cell fates during development and diseases^5–8^. Although chromatin accessibility in single-cell ATAC-seq data offers a unique opportunity to infer potential TFs bindings in heterogeneous populations^9–16^, direct measurements of TFs binding and the context of histone modifications in the same single cell have been unattainable thus far. Single-cell profiling of histone modifications and TFs binding based on the protein A/G-Tn5 transposase has been developed and widely adopted^17–21^. Recently, built on this strategy, we and others have reported a dual-omics method to simultaneously detect one epigenomic modality and transcriptome in single cells, importantly, successfully computationally linking epigenomic multimodalities via shared transcriptome as the anchor^22,23^. Recent studies reported two computational frameworks to infer the regulatory relationship between two histone modifications in single cells^24,25^. However, experimental co-assay of multiple epigenetic proteins in single cell with high throughput is relatively under-explored.

Recent advances in developing single-cell multimodal chromatin profiling methods such as scMulti-CUT&Tag^26^, MulTI-Tag^27^, nano-CT^28^ and NTT-seq^29^ allows profiling multiple histone modifications at one time in single cells. However, data sparsity, such as hundreds of reads per cell from scMulti-CUT&Tag, and up to 3,568 reads per cell in repressive histone mark H3K27me3 and hundreds of reads per cell for other active histone marks in MulTI-Tag limits its potential applications in complex biological context. Based on a single chain nanobody in fusion with Tn5, both NTT-seq and nano-CT yield higher sensitivity, however, their measurements are restricted to at most three histone modifications co-profiled at once. In addition, these technologies are not shown to be compatible for profiling TFs binding with histone modifications, nor are they used to profile epigenetic proteins and transcriptome they control in single cells.

Here, we develop a versatile strategy, dubbed CoTarget (Combined TAgmenting enRichment for multiple epiGEneTic proteins in the same cells) to jointly measure various combinations of epigenomic modalities for low-input materials and ultra-high throughput CoTarget (uCoTarget) in single cells. We show that uCoTarget is highly efficient to profile five histone modifications in single cells at the same time with high sensitivity and multimodal resolution. Notably, uCoTarget obviates tedious steps of covalent antibody-barcode conjugation and complex purification, and does not require any specialized microfluidic-based device, potentiating its wide adoptions in numerous applications. Importantly, we further develop uCoTargetX to simultaneously profile multiple epigenetic proteins and transcriptome in single cells to uncover the transcriptional outcome in the multimodal epigenomic landscapes.

## RESULTS

### Design of CoTarget for parallel profiling of multiple epigenetic proteins

We first developed CoTarget to simultaneously measure multiple histone modifications. Specific antibodies against different histone modifications or TFs were incubated with pre-assembled protein A-Tn5 with T7 barcoded adaptor (PAT-T7). The antibody-PAT-T7 complexes were guided to lightly fixed chromatin for tagmentation, and the antibody/target protein identities were distinguished by corresponding T7 barcodes and were deconvoluted in data processing. PAT-T5 complexes with secondary antibodies were then added for the further tagmentation at loci enriched for assayed epigenetic proteins. We redesigned the CoTarget library construct, making it compatible for the standard sequencing recipe (Figure 1A and Figure S1A). To achieve high sensitivity and multimodal resolution, we optimized a number of key steps in the procedure. First, we tested whether antibody-PAT T7 barcoded adaptor covalent conjugation (labeled) produce less off-target signals than non-covalent conjugation (unlabeled). In the labeled protocol, antibodies were first covalently conjugated with annealed T7 adaptors through iEDDA-click chemistry^30,31^. The antibody-T7 conjugates were then assembled with PAT to generate the stable ternary complexes. Whereas in the unlabeled group, PAT was pre-assembled with annealed T7 before incubation with antibodies without further purification (Figure 2a). Surprisingly, we found that labeled groups gave rise to markedly lower signal to noise data. The conjugates at the molar ratio 1:4.5 of antibody to adaptor even failed in target tagmentation and enrichment (Supplementary Figure 2b), suggesting that labeled antibodies may mask recognition of the epitope in histone modifications. Second, we compared the incubation of multiple antibodies, through either sequential or combined manner, with different antibody-PAT-T7 complexes. Third, we tested whether remaining PAT after the first primary antibody mediated tagmentation may induce off target effect through recruiting the subsequent primary antibody-PAT-T7 complexes through protein A domain during the second round of tagmentation. To this end, we introduced Fc proteins, to block protein A after each round of tagmentation, aiming for minimizing this undesired effect. Together, using antibodies against H3K27ac and H3K27me3 simultaneously in K562 cells, we systematically evaluated all protocols mentioned above. We compared the detection precision by calculating FRiP of each histone mark data from CoTarget experiments. We observed that the FRiP of these conditions varied from 0.55 to 0.66, among which a combination of unlabeled and sequential protocols produced the highest FRiP score, while the +Fc and no Fc protocol had modest effects on FRiP (Supplementary Figure 2c). To compare the specificity among these protocols, we made a correlation analysis. We found strong correlations between histone marks split from CoTarget data and single antibody in situ ChIP data^17^. Consistent with the FRiP analysis result, the combination of unlabeled and sequential protocols resulted in highest correlation scores with bulk ChIP-seq data, no matter +Fc or no Fc protocol was included (Supplementary Figure 2d). Thus, we adopted the combination of unlabeled, sequential and no Fc protocol for the following experiments.

**Figure 1.**
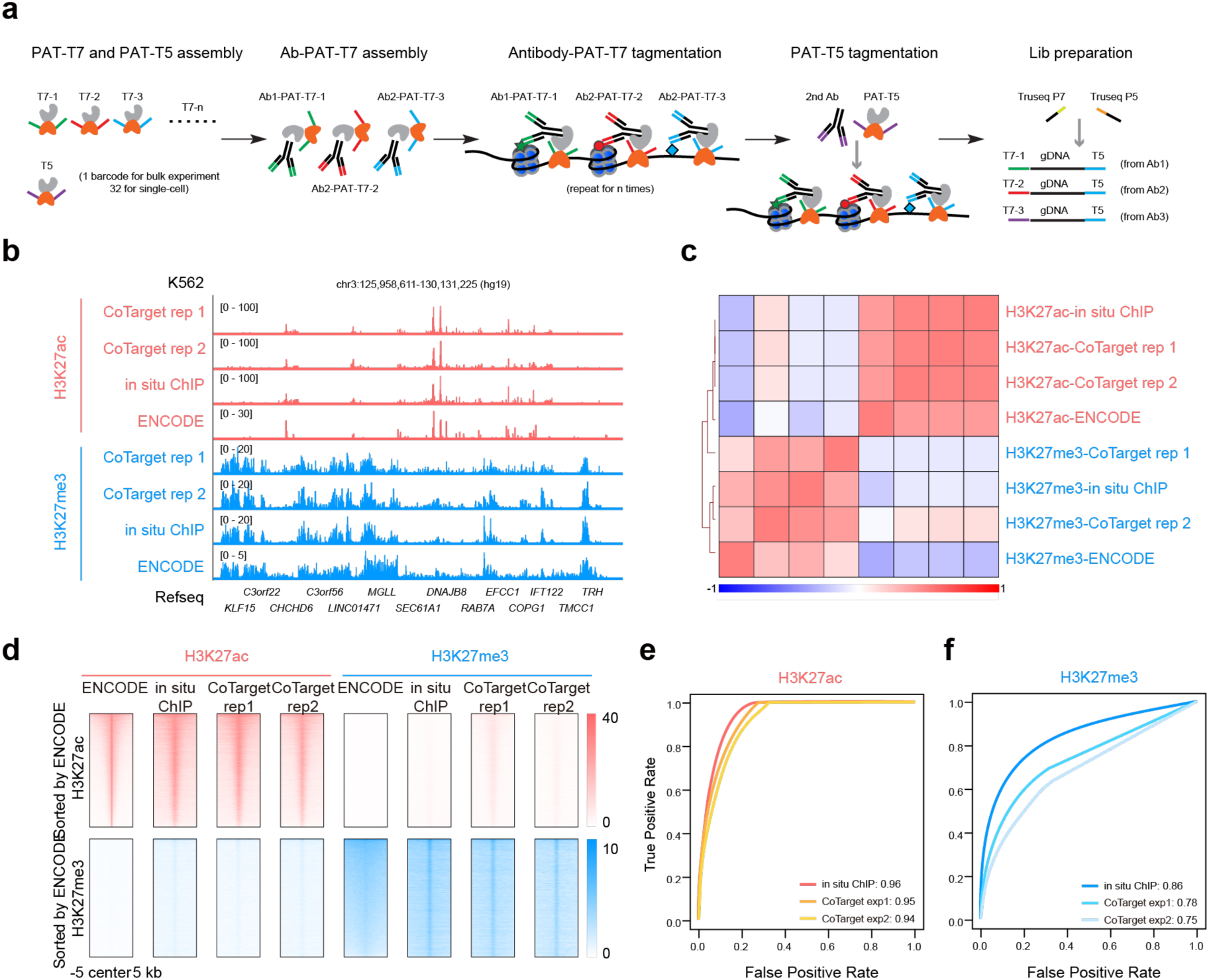
CoTarget for simultaneous profiling of multiple histone modifications. (a) Schematic of CoTarget design. (b) Track view showing H3K27ac and H3K27me3 signals at representive loci in K562 cells. (c) Heatmap showing the correlation between CoTarget, in situ ChIP, and ENCODE data in K562 cells. Hierarchical clustering of different groups was performed using the genome-wide signals in non-overlapping 5-kb windows. (d) Heatmap showing H3K27ac (up) and H3K27me3 (bottom) signals in CoTarget, in situ ChIP, and ENCODE data in K562 cells. The rows were sorted in the descending order in 31,931 ENCODE H3K27ac peaks (up) and 19,867 ENCODE H3K27me3 peaks (bottom). Bulk H3K27ac and H3K27me3 ChIP-seq data in K562 were downloaded from GSM733656 and GSM733658, respectively (e-f) Receiver operating characteristic curves for H3K27ac (e) and H3K27me3 (f) data from different groups. Values next to indicated groups are the area under the ROC curve.

**Figure 2.**
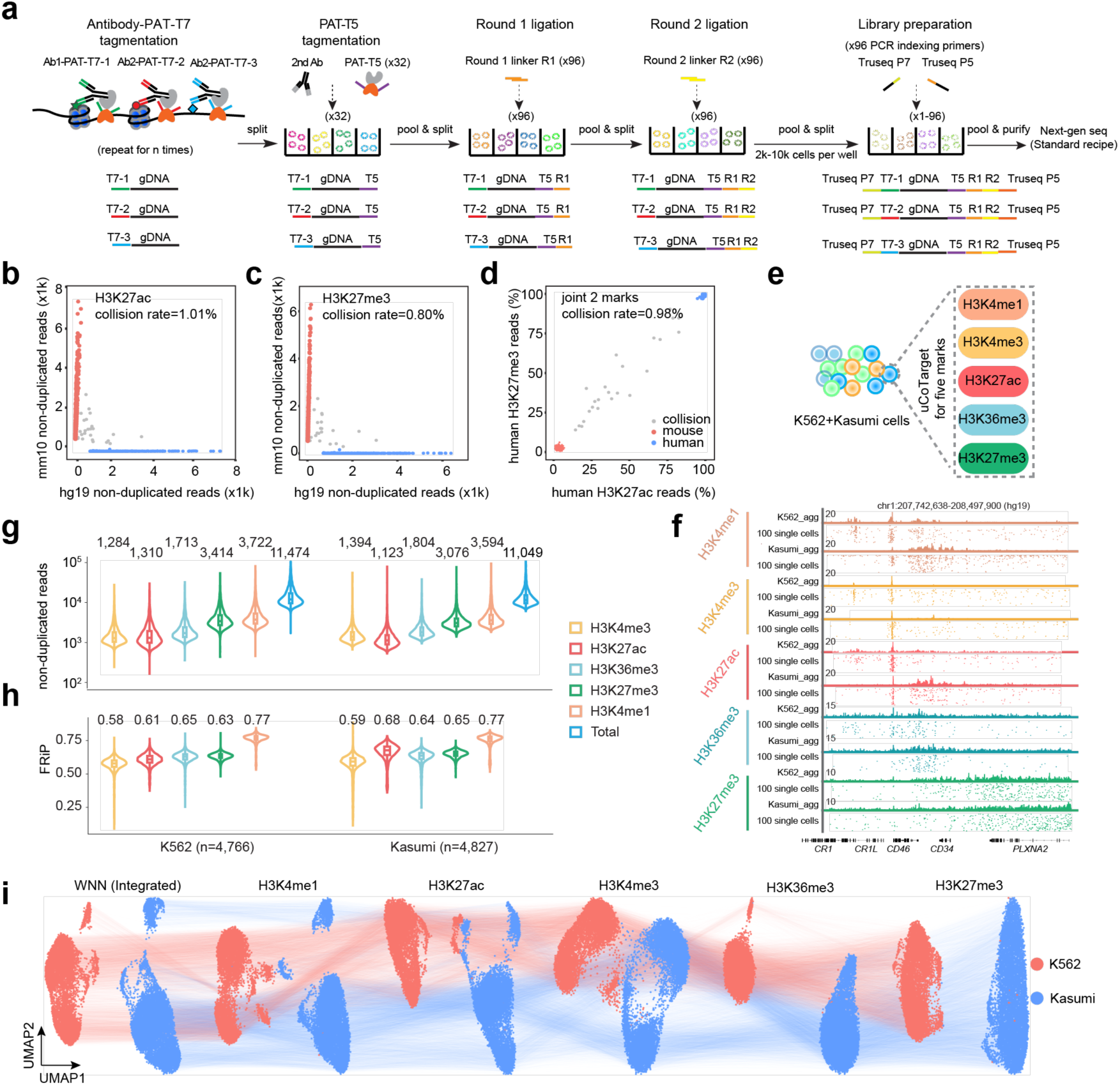
Ultra-high throughput single-cell CoTarget enables accurate profiling of five histone modifications. (a) The design of single-cell uCoTarget. (b-d) Scatter plots of the human-mouse species mixing test using H3K27ac (b), H3K27me3 (c) and joint profiles (d) of uCoTarget data in mouse ESC and human K562 cells. Points are colored by the cell identity as human (blue, >90% of reads mapping to hg 19), mouse (red, >90% of reads mapping to mm10), or collision (grey, <90% mapping to either genome). (e) Experimental design of uCoTarget profiling of five histone modifications in K562 and Kasumi-1 cells. (f) Track view showing aggregate and single-cell uCoTarget data at representive loci in K562 and Kasumi-1 cells. (g-h) Violin plots showing non-duplicated reads per cell (g) and fraction of reads in peaks (FRiP) per cell (h). Color indicates two different histone modifications.The boxes in violin plots indicate upper and lower quartiles (25th and 75th percentiles). (i) UMAP projection and visualization of single cells with integrated, H3K4me1, H3K27ac, H3K4me3, H3K36me3 and H3K27me3 information. Cells with five marks detected were used for UMAP visualization (n = 14,779).

To demonstrate the performance of CoTarget, we first applied it to K562 cells for simultaneously measuring H3K27ac and H3K27me3. Expectedly, we observed strong H3K27ac signals around representative active genes for different groups of CoTarget with 1,000 cells, *in situ* ChIP with 1,000 cells, and ENCODE ChIP-seq in K562 cells, while H3K27me3 and H3K27ac signals are mutually exclusive (Figure 1b). To further globally evaluate data quality of CoTarget, we calculated the correlation based on the signals at 5-kb bins genome wide. We found that all three datasets were highly correlated within the same histone marks, and there was a clear separation between H3K27ac and H3K27me3 datasets as shown in the correlation heatmap (Figure 1c). We next examined signals at peaks called from ENCODE ChIP-seq data as an orthogonal measurement of the performance of CoTarget. CoTarget H3K27ac and H3K27me3 data showed high enrichment at corresponding ENCODE peaks (Figure 1d). Moreover, CoTarget samples recapitulated the vast majority of H3K27ac and H3K27me3 peaks discovered in ENCODE ChIP-seq (area under curve, 0.94 and 0.95 for H3K27ac exp 1 and exp 2; 0.78 and 0.75 for H3K27me3 exp 1 and exp 2) (Figure 1e-f). Taken together, our results indicated that CoTarget enable effectively measuring multiple histone marks at once with high signal-to-noise ratio.

### Simultaneous detection of five histone modifications with ultra-high throughput single-cell CoTarget

We next established single-cell CoTarget with ultra-high throughput. To this end, we adopted the split-pool ligation-based strategy for single-cell assay^32,33^ and developed uCoTarget (ultra-high throughput CoTarget), which allowed simultaneously profiling of multiple epigenetic proteins (histone modifications and TFs) with throughput up to one million single cells in single experiment. Different antibodies were incubated with pre-assembled PAT-T7 with different T7 barcodes. Targeted tagmentation of primary antibodies was performed sequentially. Cells were then split into 32 wells for the secondary antibody-guided tagmentation with different T5 barcodes. After 2-round split-pool ligation in 96 wells, cells were combined and redistributed to 3,000-10,000 cells/well for indexed PCR as the fourth round of cell barcoding (Figure 2a, Supplementary Figure 1b). The uCoTarget design relies on 4 rounds of barcoding strategy to achieve single-cell resolution at ultra-high throughput. As a proof-of-concept, we performed H3K27ac-H3K27me3 uCoTarget species-mixing experiments in 1:1 mixed mouse embryonic stem cells (mESCs) and K562 cells. For the vast majority of cells, reads mapped either to the mouse or human genome, both for H3K27ac, H3K27me3 and joint profiles of two marks, and the cell collision rate was ∼1%, confirming a successful split-and-pool strategy of uCoTarget (Figure 2b-d). To demonstrate the performance of uCoTarget in distinguishing cell types using different epigenomic modalities, we first applied it to jointly profile H3K4me3 and H3K27me3 for the same cells in two human leukemia cell lines (K562 as chronic myeloid leukemia cells and Kasumi-1 as acute myeloid leukemia cells). We observed a nearly mutually exclusive relationship between H3K4me3 and H3K27me3 profiles from aggregate single cells (Supplementary Figure 3a-b). Aggregate profiles of K562 and Kasumi-1 cells displayed similar patterns while varied at specific peak regions, indicating the epigenetically distinct cellular states between two human leukemia cell lines (Supplementary Figure 3a). After demultiplexing and processing uCoTarget data, we obtained 7,884 single cells from an input of 8,000 cells with a median of 4,489 and 1,535 non-duplicated reads per cell for H3K27me3 and H3K4me3, respectively (Supplementary Figure 3c; Supplementary Table 3). We performed dimensionality reduction for single cells and visualized them by uniform manifold approximation and projection (UMAP) (Supplementary Figure 3d). To further demonstrate the flexibility in uCoTarget experiments, we simultaneously measured five histone modifications with two different profiling ordering (sequential H3K4me3-H3K4me1-H3K27ac-H3K36me3-H3K27me3; sequential H3K4me3-H3K4me1-H3K27ac-H3K36me3-H3K27me3) in K562 and Kasumi-1 cells (Figure 2e). The genomic profiles from aggregate and single cells displayed specific enrichment for different histone marks around representative gene loci (Figure 2f). The cell recovery rate of these uCoTarget datasets was ∼98%, yielding a median of unique reads per cell ranging from 1,123 to 3,722 and FRiP ranging from 0.58 to 0.77 with the modest sequencing depth at the duplication rate from 20.25% to 40.51% (Figure 2g-h; Supplementary Table 3). To define cell identities, we used integrated profiles as well as each histone mark profile as input for unsupervised dimensionality reduction. We found that K562 and Kasumi-1 cells were well-separated both in single modality and WNN integration (Figure 2i). Notably, epigenetic heterogeneity in cell cycle state were also identified regardless of different profiling ordering (Supplementary Figure 3g-i). Furthermore, we benchmarked uCoTarget with the most relevant other single-cell methods for joint profiling of multiple histone modifications in the same cell. In summary, uCoTarget at the modest sequencing depth manifested dramatically higher non-duplicated reads per cell and higher signal-to-noise than multi-CUT&Tag^26^, mulTI-Tag^27^, and NTT-seq^29^ (Supplementary Figure 4).

### uCoTarget for profiling of three marks during generation of HSPC

Next, we used uCoTarget to investigate how the different epigenomic modalities coordinates cell lineage specification using the model of hematopoietic stem/progenitor cells (HSPCs) differentiation. We harvested cells at differentiation day 8 starting from human embryonic stem cells (hESCs) based on a chemically defined hematopoietic differentiation protocol^34^, and performed uCoTarget experiments for simultaneously detecting H3K27me3, H3K27ac, and H3K4me3 in the same cells (Figure 3a; Supplementary Figure 5a). After filtering out low-quality cells and potential doublets from two sublibraries, we obtained 14,002 cells with a median of 1,851, 11,437, and 6,078 unique reads per cell for H3K4me3, H3K27ac and H3K27me3, respectively (Supplementary Figure 5b-d). Clustering single cells by H3K27ac profiles identified three major populations, annotated as mesoderm, hematopoietic cells, and cardiac cells (Figure 3b). We excluded cardiac cells to focus on the HSPC differentiation (Figure 3c; Supplementary Figure 5e). ChromVAR analysis^15^uncovered dynamic changes of a set of core transcription factors (TFs) from mesoderm cells to HSPCs, and also validated the identification of HSPC differentiation path (Supplementary Figure 5f). To investigate the possibility of different histone marks in deconvolution of cell types, we performed dimensionality reduction analysis using the H3K4me3, H3K27me3 and H3K27ac profiles separately. Not surprisingly, H3K27ac outperformed other marks for de novo cell type identification, supporting the lineage-specifying role of enhancers in the context of HSPC differentiation (Figure 3d). We next examined the single-cell kinetics of different histone marks during the HSPC differentiation. SMAD3 and TAL1 are known important regulators for mesodermal and blood development, respectively^35,36^. Expectedly, we observed strong H3K4me3 and H3K27ac signals in mesoderm cells at *SMAD3* loci and in HSPCs at *TAL1* loci (Figure 3e-f). Interestingly, H3K27me3 signals around *TAL1* in mesoderm and cardiac cells likely repressed its expression in accordance with lower H3K27ac and H3K4me3 (Figure 3e-f).

**Figure 3.**
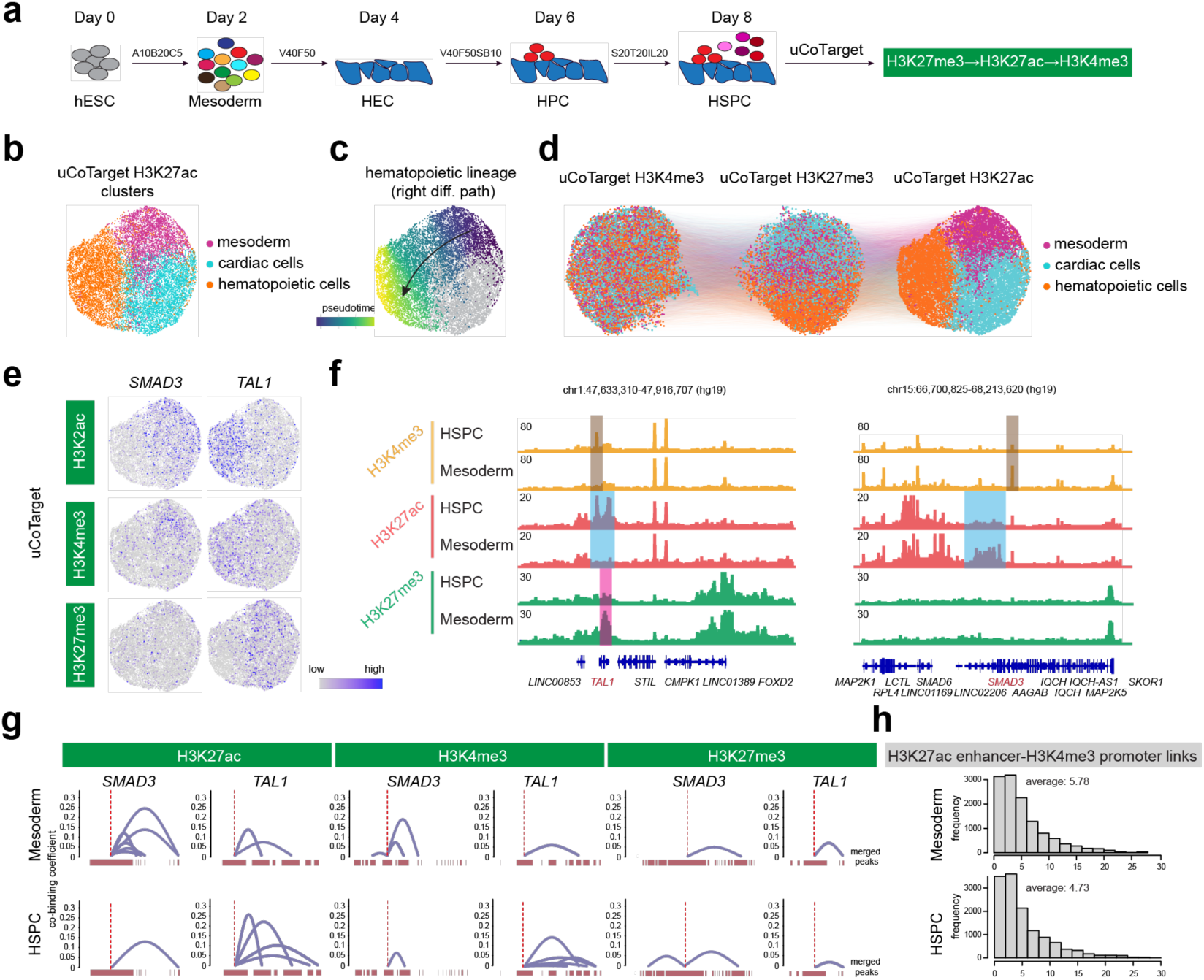
uCoTarget profiling of histone modifications during generation of haematopietic stem progenitor cells. (a) Workflow of human ESC differentiation towards HSPCs. At differentiation day 8, cells were digested to single cells and collected for H3K27me3-H3K27ac-H3K4me3 uCoTarget experiments. hESC, human embryonic stem cell. HEC, hemogenic endothelial cell. HPC, hematopoietic progenitor cell. HSPC, hematopoietic stem and progenitor cell. (b) UMAP showing three major subpopulations identified by uCoTarget-H3K27ac data during generation of HSPCs. Three clusters were identified and annotated, n = 3,273, 5,365, 5,364 for mesoderm, hematopoietic cells, and cardiac cells, respectively. (c) Pseudotime analysis of hematopoietic lineage. The color from darkblue to yellow indicates pseudotime from early to late. For hematopoietic lineages, cardiac cells colored by grey were not used for pseudotime analysis. (d) UMAP showing single cells with integrated and connected information of three histone modifications. (e) Projection of uCoTarget-H3K27ac, uCoTarget-H3K4me3, and uCoTarget-H3K27me3 around representative loci. The color from grey to purple indicates signals from low to high. (f) Track view showing the aggregate signals of HSPC and mesoderm cells for three histone marks around TAL1 (left) and SMAD3 (right). (g) Co-binding analysis of mesoderm cells (top) and HSPC (bottom) based on signals of uCoTarget-H3K27ac, uCoTarget-H3K4me3, and uCoTarget-H3K27me3 by Cicero. The linkages identified by Cicero were visualized around the genomic regions of “chr15:66970886-67874328”, and “chr1:47482049, 48022416” for SMAD3 and TAL1, respectively. (h) Distribution of numbers of H3K27ac enhancer-H3K4me3 promoter links around TSS ± 500 kb of genes.

To further explore potential interactions of cis-regulatory elements coordinating cell-fate regulating gene activation, we used Cicero ^37^ to predict co-binding sites for mesodermal cells and HSPCs. *SMAD3* enhancers have strong putative co-binding signals in mesodermal cells, while remarkably lower in HSPCs. H3K4me3 also exhibited cell-type specific co-binding events while the co-binding coefficient by H3K4me3 was lower than that of H3K27ac, indicating that co-binding of cell-type specific enhancers more likely drive gene expression specificity (Figure 3g). Notably, uCoTarget enabled us to jointly analyze multifactor defined cis-regulatory elements. We found that the average number of co-binding events identified by H3K27ac distal enhancer-H3K4me3 promoter regulating corresponding target genes in mesoderm cells was significantly higher than that in HSPC at genes ± 500-kb regions (P=8.4×10−5, Mann–Whitney U-test; Figure 3h). Together, our results support the importance of cell-type specific elucidation of multimodal epigenetic regulation of HSPC differentiation.

### RUNX1 chromatin engagement priming for H3K27ac along the HSPC path

TFs play an instructive role in coordinating multimodal epigenomic dynamics and regulating cell fate specification^38^. We performed H3K27ac-RUNX1-H3K27me3 uCoTarget experiments to explore how different modalities change in shaping the epigenomic landscape during the HSPC differentiation (Figure 4a). RUNX1 is a master TF regulating hematopoiesis^39–41^. It has been reported that RUNX1 regulates GFI1 and GFI1B to promote the hemogenic endothelial program^42^. We found that strong RUNX1 enrichment accompanying H3K27ac were observed around GFI1B, while lower RUNX1 and strong repressive histone mark H3K27me3 at GFI1 loci (Figure 4b). We next asked how RUNX1 coordinates with H3K27ac and H3K27me3 during HSPC differentiation at single-cell level. We calculated Cramér’s V^27^ to quantify the association of RUNX1-H3K27ac, RUNX1-H3K27me3 in single cells. The Cramér’s V in RUNX1-H3K27ac were ∼3-fold higher than that in RUNX1-H3K27me3, indicating a more profound interplay between RUNX1 and H3K27ac in setting up the active epigenome while we did not exclude the repressive role of RUNX1 on co-targeting with H3K27me3 (Figure 4c). Furthermore, we identified major clusters and HSPC differentiation trajectory in accordance with the results from uCoTarget in profiling three histone modifications as above (Figure 3b-d, Figure 4d). To examine the dynamic change of RUNX1 and histone modifications during transition from mesoderm cells to HSPCs, we performed differential analysis on uCoTarget-H3K27ac data. 429 differential enhancer regions including SMAD3, TEAD2 (mesoderm-specific), ERG and TAL1 (HSPC-specific) were identified (Figure 4e). The dynamic trend in both RUNX1 and H3K27ac signals at the corresponding enhancer-linked regions was well correlated along pseudotime, while lower H3K27me3 was accompanied (Figure 4f). Interestingly, we found that RUNX1 binding preceded H3K27ac activation at HSPC-specific enhancer regions during the generation of HSPCs (Figure 4f). RNA Velocity has been widely used to predict the future state of individual cells by estimating the ratio of rates of gene splicing and degradation via the relative abundance of unspliced and spliced mRNAs^43,44^. Based on the similar relationship of time-dependent processes between RUNX1/H3K27ac and unspliced/spliced mRNAs, we performed chromatin velocity-based trajectory inference as previously described^28,45^ in the context of HSPC differentiation using uCoTarget-RUNX1 data as input into the unspliced layer and uCoTarget-H3K27ac data into the spliced layer. We found that chromatin velocity largely recapitulated the dynamics of HSPC differentiation (Figure 4g). Together, these results demonstrated that RUNX1 priming for H3K27ac orchestrates the emerging epigenomic landscape during the generation of HSPCs.

**Figure 4.**
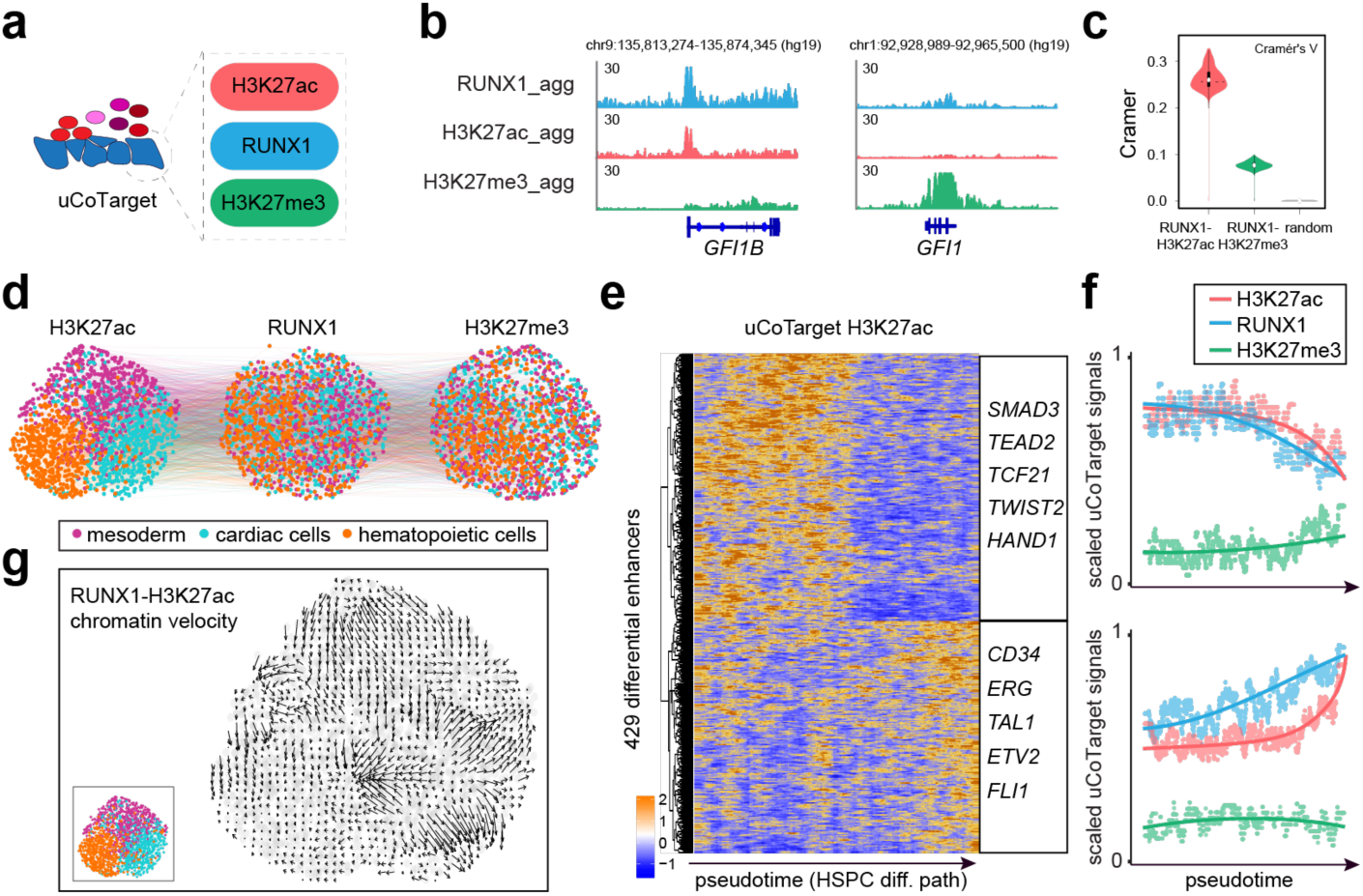
uCoTarget reveals that RUNX1 priming H3K27ac orchestrates the emergence of haematopietic stem progenitor cells. (a) uCoTarget profiling of human ESX differentiation toward HSPCs for transcription factor RUNX1 and histone modifica­tions. (b) Track view showing aggregate RUNX1, H3K27ac, and H3K27me3 signals at representive loci. (c) Violin plots showing calculated Cramer’s V of association between different target combinations. (d) UMAP showing single cells with H3K27ac (left), RUNX1 (middle), and H3K27me3 (right). Single cells were connected based on the modality information from the same cells (n = 1,786). (e) Heatmap showing uCoTarget H3K27ac signals at 429 differential enhancers (Wilcoxon test, log fold change > 0.25, P<0.01) along HSPC differentiation pseudotime. Representative gene loci are listed alongside the heatmap. (f) Line plots showing average RUNX1, H3K27ac, and H3K27me3 signals at two enhancer clusters as shown in **e.** The y-axis is the scaled uCoTarget region score for each modality, and the x-axis reprents pseudotime of HSPC differentiation trajectory. The line depicts the calculated smoothed signals (rolling window = 20, step =1) followed by local polynomial regression fit (loess) of uCoTarget data for H3K27ac (red), RUNX1 (blue), and H3K27me3 (green). (g) Chromatin velocity analysis based on the UMAP in d. The chromatin velocity was calculated using uCoTarget-RUNX1 data as input into the unspliced layer and uCoTarget-H3K27ac data into the spliced layer and then running velocyto.R (https://github.eom/velocyto-team/velocyto.R) with default parameters. The length of the arrows indicate the chromatin velocity and the direction of the arrows represent the inferred future state of the position.

### UCoTargetX enables single-cell co-assay of transcriptome and multimodal epigenome

Next, we attempted to jointly measure multimodal epigenome and transcriptome based on uCoTarget to uncover the epigenetic regulation of transcription. We successfully developed a method, dubbed uCoTargetX, by combining the uCoTarget workflow with a modified Smart-seq2 procedure to obtain the multimodal readout of polyadenylated messenger RNA and multiple chromatin modalities (Figure 5a). We first evaluated the potential effect of the order of DNA or RNA modality barcoding by performing RNA-H3K27ac-H3K27me3 and H3K27ac-H3K27me3-RNA uCoTargetX experiments in HEK 293 cells. We found that slightly higher unique reads and genes per cell were observed in the condition of barcoding RNA first (Supplementary Figure 6). Thereafter, we further applied this condition to jointly compare gene expression and enrichment of histone modifications between H1 and K562 cells. Expectedly, highly expressed genes were associated with high H3K27ac and low H3K27me3 and vice versa (Figure 5b). We identified a median yield ranging from 5,507 to 26,705 unique reads with joint capture of 1,000∼2,000 genes per cell in both H1 and K562 cells (Figure 5c-e). We then constructed cell-by-bin matrix for each epigenomic modality and cell-by-gene matrix for RNA modality, and performed dimensionality reduction to visualize individual cells. K562 and H1 cells were well-separated using each modality and integrated WNN (Figure 5f). In addition, RNA modality improved the accuracy of cell type identification by analyzing the proportion of neighbors belonging to the same cell type using a single modality or different combination of modalities (Figure 5g). To benchmark uCoTargetX in a more dynamic biological context, we performed RNA-RUNX1-H3K27ac uCoTargetX experiments in CD34+ and CD34-cells during the HSPC differentiation (Figure 5h). Differential enrichments for H3K27ac and RUNX1 binding were observed around *KAT6B* gene locus while not seen at the RNA level, indicating the asynchronous change between epigenome and transcriptome (Figure 5i). Integrated UMAP exhibited a differentiation trajectory from CD34- to CD34+ cells, also validating the data quality of multimodal profiling (Figure 5j). We further calculated Cramer’ V to evaluate the association between transcriptome and DNA modalities. RNA modality was more associated with H3K27ac than RUNX1 modality for both CD34+ and CD34- cells (Figure 5k).

**Figure 5.**
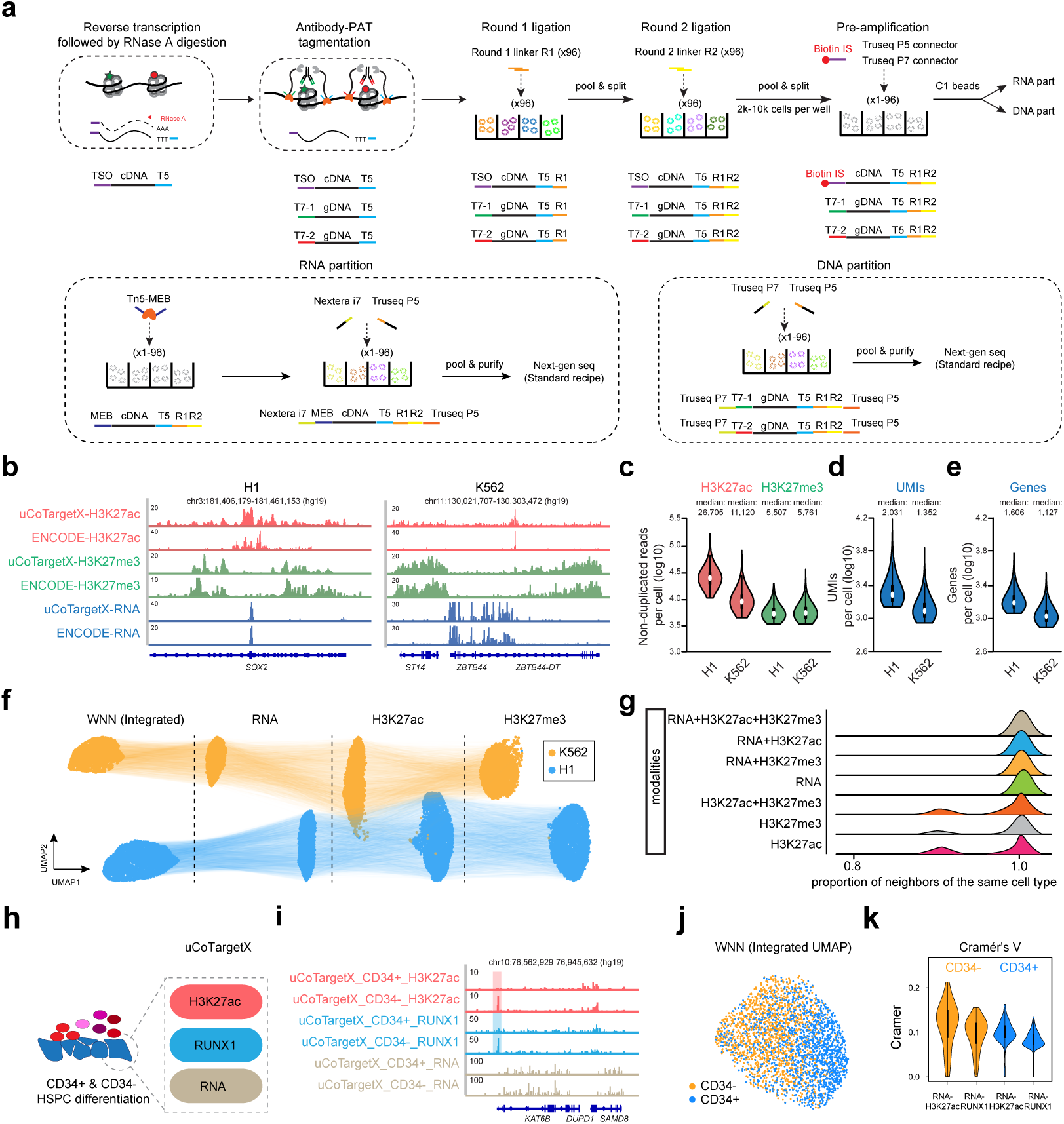
uCoTargetX enables joint detection of transcriptome and multiple histone modifications in single cells. (a) Schematic of uCoTargetX experimental design. (b) Track view displaying both RNA, H3K27ac and H3K27me3 signals on representative gene loci in K562 and H1 cells. (c-e) Violin plots showing non-duplicated reads per cell (c), detected UMI (d), and genes (e) per cell. The boxes in violin plots indicate upper and lower quartiles (25th and 75th percentiles). (f) UMAP projection and visualization of single cells with integrated, RNA, H3K27ac and H3K27me3 information. Cells with each modality detected were used for UMAP visualization. (g) Proportion of neighbors belonging to the same cell type using a single modality or different combination of modalities. (h) uCoTargetX profiling of human ESC differentiation toward HSPCs for transcription factor RUNX1, H3K27ac and RNA. (i) Track view displaying both H3K27ac, RUNX1 and RNA signals on representative gene loci for CD34+ and CD34- cells during the generation of HSPC. (j) UMAP projection and visualization of single cells during the generation of HSPCs. (k) Violin plots showing calculated Cramer’s V of association between different target combinations.

## DISCUSSION

In sum, we present a generalizable, versatile method for simultaneous measurements of multiple epigenetic modalities with ultra-high throughput. The uCoTarget design makes it compatible with the standard Illumina sequencing platform without need of specialized microfluidic device, and uCoTarget can be easily implemented with the extremely lower cost of $ 0.01 per cell in major biochemical reagents. Notably, the throughput in uCoTarget can be easily scaled up to 1 million single cells by simply sequencing the sublibraries from one 96-well PCR plate. Importantly, uCoTarget enables joint measurement of five histone modifications at the same time. We believe they are readily expanded to more epigenomic modalities since chromatin protein complexes onto DNA are slightly fixed, potentially allowing for many rounds of antibody incubation-tagmentation. We showcase that uCoTarget enables exploring the single-cell dynamics of multimodal epigenomic landscape during the HSPC differentiation in profiling of histone modifications and TFs. Our data demonstrate that RUNX1 precedes H3K27ac in setting up the HSPC-specific epigenome. Importantly, uCoTarget can be easily upgraded to uCoTargetX to measure RNA together with multiple histone modifications simultaneously, in aim to analyze multimodal heterogeneities and uncover associations between different modalities.

Compared to recently reported single-cell multifactorial epigenomic technologies, uCoTarget is more convenient and efficient to achieve multi-modality profiling. Both nano-CT and NTT-seq relies on fusion protein nanobody-Tn5, limited by the available different species-raised primary antibodies in matching corresponding single chain nanobodies for at most three epigenetic proteins co-profiled in one experiment to date. Further, compared with nano-CT and NTT-seq, uCoTarget does not require production of new different recombinant nanobody–Tn5 fusion proteins, is compatible with any commercial antibodies for simultaneous measurements, and enable easy co-profiling of 5 to more epigenetic proteins in single experiment with ultra-high throughput. In order to remove uncomplexed antibodies and free adapters, multi-CUT&TAG used TALON beads for affinity-purification before each experiment, which was essential for them to partition reads with specific barcodes to each antibody to each epigenomic modality. On the contrary, uCoTarget obviates any purification step since we attest that extensive wash in conjunction with slight cell fixation help remove excess free adaptors, which otherwise induce cross-modality contaminations. MulTI-Tag took on the strategy to covalently conjugate PAT-T5 adaptors with primary antibodies. However, after thorough evaluation of various conditions, we find an optimal combination including non-conjugate antibody (unlabeled) and sequential antibody incubation (Supplementary Figure 2). Instead, we demonstrate that covalent antibody-barcode conjugation is prone to mask the recognition of the epitope in histone modifications or other epigenetic proteins. With a unique feature of co-assay of transcriptome and multiple chromatin profiles in this work, uCoTargetX lends itself to become the most convenient and easy-to-use single-cell multifactorial epigenomic profiling tool (Supplementary Figure 4a,c).

Nevertheless, current single-cell epigenomic and multimodal approaches are far from mature due to the data sparsity^46,47^. We obtained up to 19,366 reads per cell in the summed multimodality in uCoTarget, suggesting that optimization in molecular biology holds promise to further improve the sensitivity. The higher dimensional reconstruction of the genome-epigenome-transcriptome-proteome landscape in single cells would provide new insights into regulatory mechanisms in cell fate specification^62–65^. We also foresee that uCoTargetX holds the potential to be applied in constructing human cell atlases and uncovers unbiased epigenetic principles in deregulated cell lineages in human diseases.

## ACCESSION CODE

Raw sequencing data have been deposited at the NCBI Gene Expression Omnibus (GEO) with the accession number GSE220193 (reviewer token: yrozmekuzzenpwp), https://www.ncbi.nlm.nih.gov/geo/query/acc.cgi?&acc=GSE220193.

## Acknowledgements

We thank all members of the He lab for critical comments on this manuscript. Part of the analyses was performed on the High Performance Computing Platform of the Center for Life Sciences, Peking University. We thank the flow cytometry Core at National Center for Protein Sciences at Peking University, particularly Liying Du and Huan Yang, for technical help. A.H. was supported by the National Key Research and Development Program of China (2021YFA1100100 and 2019YFA0801802), the National Natural Science Foundation of China (32192401 and 32025015), the Peking-Tsinghua Center for Life Sciences, and the 1000 Youth Talents Program of China.

## Author contributions

A.H. conceived and designed the study. Q.W. designed and performed all experiments, assisted with C.C.L., H.X. performed the computational analyses, supervised by A.H., and H.X., Q.W. and A.H. wrote the paper with input from all other authors. All participated in data discussion and interpretation.

## Competing interests

The authors declare no competing interests.

## Cell culture

Wild-type V6.5 murine embryonic stem cells (mESCs) were cultured on 0.1% gel-atin-coated plates in ESC DMEM culture medium containing 15% fetal bovine serum (Sigma), 1% Penicillin/Streptomycin (Hyclone), 1% Glutamax (Hyclone), 0.1 mM 2-mercaptoethanol (Sigma), 1% MEM nonessential amino acids (Cellgro), 1% nucleoside (Millipore), and 1,000 U/ml recombinant leukemia inhibitory factor (LIF) (Millipore). V6.5 cells were digested with 0.1% Trypsin (Hyclone) at 37°C for 3 min to harvest single cells. K562 and Kasumi-1 cells were cultured in RPMI-1640 medium containing 10% fetal bovine serum (Sigma), 1% Penicillin/Streptomycin (Hyclone) and 1% Glutamax (Hyclone). K562 and Kasumi-1 cells were harvested by centrifugation for 2 min at 600 g. H1 human embryonic stem cells (hESCs) were cultured in matrigel (Corning)-coated plates using hESC culture medium mTeSR1. H1 cells were digested with Accutase (Thermo) at 37°C for 3 min to harvest single cells.

## HSPC differentiation

HSPCs (hematopoietic stem and progenitor cells) were differentiated from H1 hESCs as previously reported^1^. In short, dissociated hESCs were placed into several wells of 12-well plate at ∼50k cells/well. hESCs were then induced under the step-by-step differentiation condition after overnight growth. During the first 2 days, 10 ng/ml Activin A, 20 ng/ml BMP4 and 5 μM CHIR99021 were added to the basic RPMI-1640 medium containing 1% Glutamax (Hyclone), 1% MEM nonessential amino acids (Cellgro), 1% Penicillin/Streptomycin (Hyclone), 0.1 mM 2-mercaptoethanol (Sigma), 2% B27 and 1% insulin-transferrin-selenium (ITS; Gibco), which induced hESCs transition into early mesoderm stage. Second, 40 ng/ml VEGF, and 50 ng/ml bFGF were added in the basic medium for the other 2-day induction, during which cells tended to be HE (hemogenic endothelium). Third, 40 ng/ml VEGF, 50 ng/ml bFGF, and 10 μM SB431542 were added to the basic medium from day 4 to day 6 to turn cell fate to HPC (hemogenic progenitor cells). From day 6 to day 8, cells are treated with 20 ng/ml SCF, 20 ng/ml TPO, and 20 ng/ml IL-3 in basic IMDM medium, resulting into formation of HSPCs and mature hemogenic cells. At day 8, differentiated cells were digested with Accutase (Thermo) at 37°C for 3 min to harvest single cell suspension.

## Sample fixation (Formaldehyde and methanol)

Dissociated cells were washed once with 0.1% BSA-PBS and fixed by 0.25% formaldehyde (FA) on ice for 5-10 min (more fragile cells may require longer fixation time in pilot experiment test). The reaction was quenched by adding 35 mM glycine on ice for 5 min. The cells were washed twice with 0.1% BSA-PBS and resuspended with 100 μl 0.1% BSA-PBS. 900 μl pre-chilled -20°C methanol was added dropwise. The fixed samples can be stored at -80°C for at least one year.

## Antibodies used in this study

Antibodies used in this study were listed: H3K27me3 (Active motif, Cat: 39155, Lot: 19610006), H3K27ac (Abcam, Cat: ab4729, Lot: GR3305164-1), H3K36me3 (Active motif, Cat: 61101, Lot: 28818005), H3K4me3 (Millipore, Cat: 04-745, Lot: 3543820), H3K4me1 (Abcam, Cat: ab8895-50, Lot: 768604), and RUNX1 (Abcam, Cat: ab23980, Lot: GR3364239-1).

## Barcoded protein A-Tn5 preparation

Protein A-Tn5 (PAT) expression, purification, and assembly were performed as previously described^2^. In brief, pET28a-His-pA-Tn5 expression vector was transformed in chemically competent Bl21 (DE3). One single clone was inoculated into 100 ml LB medium and grew overnight. 10 ml over-night growing cells were transferred into 1 L LB medium and the culture was incubated at 220 rpm, 37°C for 3-4 hours until the O.D. reaching 0.8. The culture was fully chilled on ice for 30 min, IPTG was added to the culture at 0.2 mM and the culture was further incubated at 23°C, 80 rpm for 5 hours for PAT expression. Bacterial pellets were collected and lysed in HXG buffer (20 mM HEPES-KOH, pH 7.2, 0.8 M NaCl, 10% glycerol, 0.2% Triton X-100). PAT was released by sonication and bacterial genomic DNA was fully removed by PEI (Sigma P3143). PAT was then purified by Ni-NTA column (QIAGEN). The elution from Ni-NTA column was dialyzed overnight to remove imidazole. At last, PAT was concentrated using a 30 kDa cutoff ultracentrifuge column (Millipore) and stored at -20°C before use. The quantity of purified PAT was visualized by Coomassie bright blue staining after PAT was resolved on 7.5% SDS-PAGE. PAT and annealed barcoded adaptors (Supplementary table 2) were mixed at an equal molarity at 37.5 μM. The sample was incubated at 25°C for 60 min and then transferred to -20°C before use. The assembled PAT is stable and may be stored at -20°C for 1-2 years.

## Antibody-PAT-T7 complex assembly

0.5 μg antibody (3.33 pmol), 0.22 μl 37.5 μM pre-assembled T7 barcoded PAT (8.25 pmol), and 5 μl Wash buffer (1 ml 1 M HEPES pH 7.5, 1.5 ml 5 M NaCl, 12.5 μl 2 M spermidine, 10 mM sodium butyrate, and the final volume to 50 ml with ddH_2_O) were mixed thoroughly and incubated at room temperature for 1 hour. The Antibody-PAT-T7 complex is suggested to be used in following experiments within one day of assembly. We also compared our strategy (named unlabeled group) with recently published MulTI-Tag^3^, which achieved multifactorial profiling through antibody-oligo conjugation (covalent conjugation, named labeled group). For labeled group, 0.5 μg T7-antibody conjugates (3.33 pmol), 1 μl 5 μM PAT (5 pmol), and 16 pmol unconjugated annealed T7 adaptor of the same sequence were mixed thoroughly and incubated at room temperature for 1 hour.

## Antibody oligo covalent conjugation

The conjugation procedure was only required for the labeled group. Antibodies and oligonucleotides were covalently and irreversibly conjugated by iEDDA-click chemistry as previously described^4,5^ with a few changes.

### Oligo labeling

5’-NH2-C12 modified T7 (5’-[NH2-C12] GTCTCGTGGGCTCGGCTGTCCCTGTCC[6 bp Ab barcode]AGATGTGTATAA-GAGACAG) and 5’-Phos-Tn5MErev (5’-[Phos] CTGTCTCTTATACACATCT) were dissolved with 1×PBS to 200 μM and annealed at equal molar ratio to generate 100 μM T7 adaptors. 30 μl 100 μM annealed T7 adaptor were mixed with 4.5 μl 10×BBS, 4.5 μl DMSO and 6 μl 10 mM TCO-PEG4-NHS (Click Chemistry Tools) and the sample was incubated at room temperature for 30 min. 2.5 μl 50 mM glycine pH 8.5 was then added to quench residual NHS groups. TCO labeled T7 adaptor was desalted using Micro Bio-Spin P-6 columns (Bio-Rad). The concentration of labeled T7 adaptor was quantified. TCO labeled T7 adaptors may be stored at 4°C before use for no longer than 1 week.

### Antibody labeling

Antibodies without carrier proteins (e.g. BSA or gelatin) can be labeled directly, and antibodies with carriers need to be purified before subjected to labeling below. 15 μg antibody was washed twice with 450 μl 1×BBS by ultra-centrifugation with an Ultra-0.5 30 kDa MWCO filter unit (Millipore). The ultra-centrifugation was performed at 14,000 g for 5-10 min each time until the sample reached its minimal volume (∼15 μl). The antibody was then transferred to a new tube and mixed with 0.5 μl 2 mM mTz-PEG4-NHS (Click Chemistry Tools) (the molar ratio of antibody to mTz-PEG4-NHS is 1:10) and incubated at room temperature for 30 min. 2 μl 50 mM glycine pH 8.5 was then added to quench any unreacted NHS groups. mTz labeled antibody was washed twice with 450 μl 1×BBS as described above and carefully transferred to a new tube for reaction with labeled adaptors.

### Conjugation

15 μg mTz labeled antibody (100 pmol) was mixed with 10-150 pmol TCO labeled T7 adaptors at minimal volume and the sample was incubated at 4°C overnight. The molar ratio of antibody: adaptor heavily affects the degree of conjugation, which obeys the Poisson distribution. After the reaction was complete, 1/10 of the reaction volume of 10 mM TCO-PEG4-gly was added to quench any residual tetrazine reaction sites on the antibody. Sodium azide was added to 0.02% (m/v) for long-term storage. The labeled sample may be stored at 4°C for half a year until use. The conjugation degree can be visualized by non-reducing SDS-PAGE analysis.

### uCoTarget

#### Preparation of oligonucleotides for ligations

All oligos used for ligation (Supplementary Table 2) were dissolved with STE buffer (10 mM Tris-HCl pH 8.0, 50 mM NaCl, and 1mM EDTA), round 1 linker to 90 µM, round 1 barcodes to 100 µM, round 2 linker to 110 µM, and round 2 barcodes to 120 µM. Equal volume of round 1 linker and round 1 barcodes were mixed and annealed to generate hybridization stock round 1 containing 45 µM linker and 50 µM barcodes. Equal volume of round 2 linker and round 2 barcodes were mixed and annealed to generate hybridization stock round 2 containing 55 µM linker and 60 µM barcodes. Hybridization stock round 1 and round 2 were then diluted 20-fold with STE buffer to working solution, named round 1 adaptors and round 2 adaptors, respectively. Stocking solution was stored at -20°C and was re-annealed after each cyle of freeze and thawing. The working solution was stored at 4°C and suggested to be used within 1 month.

### Antibody-PAT-T7 targeting tagmentation

Two distinct strategies were designed for targeting tagmentation. For sequential strategy, fixed cells were washed twice with 0.1% BSA-PBS for rehydration, and incubated with the first Antibody-PAT-T7 complex in 100 μl Wash-buffer-TX-high salt (Wash buffer with 0.01% Digitonin, 0.05% TX-100, and 300 mM NaCl) plus 2 mM EDTA in room temperature for 1 hour. After incubation, cells were washed 3 times with Wash-buffer-TX-high salt to remove free unbound Antibody-PAT-T7 complex and resuspended with 50 μl Reaction buffer (10 mM TAPS pH 8.3, 10 mM MgCl_2_, 10 mM sodium butyrate, 0.01% Digitonin). The reaction was incubated at 37°C for 1 hour at a thermal cycler. Hot lid is set at 40°C. The reaction was stopped by incubating cells in 180 µl Wash-buffer-TX-high salt with 5 mM EDTA at room temperature for 5 min. The above steps were repeated N times for the other Antibody-PAT-T7 complex. For combined strategy, all Antibody-PAT-T7 complexes were added to cells at once and targeting tagmentation reactions were performed simultaneously. After careful evaluation of these strategies, we adopted the sequential manner in following single-cell experiments.

### Secondary antibody guided PAT-T5 tagmentation (1st round barcoding by PAT-T5)

After targeting tagmentation, cells were incubated with secondary antibodies at 1:1000 dilution in 100 μl Wash-buffer-TX (Wash buffer adding 0.01% Digitonin, 0.05% TX-100) at room temperature for 1 hour. Unbound secondary antibodies were removed by washing cells twice with Wash-buffer-Dig (Wash buffer adding 0.01% Digitonin). Cells were split into 32 wells with different PAT-T5 (Supplementary Table 2) (37.5 µM PAT-T5 was diluted at 450-fold in the reaction) in each well in 100 μl Wash-buffer-TX-high salt. The reaction was performed at room temperature for 1 hour. Unbound PAT-T5 was removed by washing cells twice with Wash-buffer-TX-high salt. Cells were resuspended with 50 μl reaction buffer and incubated at 37°C for 1 hour at a thermal cycler. Hot lid is set at 40°C. The reaction was stopped by incubating cells in 180 µl Wash-buffer-TX-high salt with 5 mM EDTA at room temperature for 5 min. All cells were combined and washed twice with NSB (nuclei suspension buffer) (10 mM Tris-HCl, pH 7.5, 10 mM NaCl, 3mM MgCl_2_, and 0.01% TX-100).

### 2-round ligation (second to third round barcoding by ligation)

Cells were resuspended with 1 ml NSB and distributed to 96 wells at 10 µl each well with 40 µl 1st round ligation mix, comprising 5 µl 10×T4 ligation buffer (NEB), 0.2 µl 10% TX-100, 10 µl NSB, 22.8 µl ddH_2_O, and 2 µl round 1 adaptor. The reaction was incubated at room temperature for 30 min with gentle shaking (300 rpm). 10 µl round 1 blocking mix was added to each well, which contains 0.11 µl 100 µM round 1 blocking, 2 µl 10×T4 ligation buffer, and 7.89 µl ddH_2_O. The reaction was incubated at room temperature for another 30 min with gentle shaking. All cells from 96 wells were combined and distributed to a new plate of 96 wells at 50 µl each well with 10 µl round 2 hybridization mix in each well, comprising 2 µl round 2 adaptor and 8 µl ddH_2_O. The reaction was incubated at room temperature for another 30 min with gentle shaking. 10 µl round 2 blocking mix was added to each well containing 0.132 µl 100 µM round 2 blocking, 1 µl 1% TX-100, and 8.868 µl ddH_2_O. The reaction was incubated at room temperature for another 30 min with gentle shaking. All cells from 96 wells were combined, washed twice with NSB, and resuspended in 200 µl final ligation solution, consisting of 20 µl 10×T4 ligation buffer, 10 µl 400 U/µl T4 Ligase (NEB), 10 µl 1% TX-100, 40 µl NSB, and 120 µl ddH_2_O. Final ligation was performed at room temperature for 30 min with gentle shaking. Final ligation can be scaled down to 20 µl for each sample in pilot experiments. Cells were washed twice with NSB and cell numbers were counted with hemocytometer. Cells were diluted with 0.1% BSA-PBS to 3,000-5,000/µl and were aliquoted to 96-well plates as many as possible at 1 µl/well. Each well was pre-set with 4 µl lysis buffer (10 mM Tris-HCl pH 8.5, 0.05% SDS and 0.1 mg/ml Proteinase K). The mix was incubated at 55°C for 30 min for cell lysis. 1 µl 1.8% TX-100 and 1 µl 10 mM PMSF (Sigma) was added to each well and the reaction was incubated at 37°C for 15 min to quench SDS and deactivate proteinase K.

### 2-round PCR for library preparation (fourth round barcoding by PCR)

To make the resulting libraries fit to the standard Illumina sequencing recipe, we adapted two rounds of PCR enrichment. 1st round PCR mix was prepared as follows: 5 µl cell lysates, 6 µl 5xHiFi buffer, 0.9 µl 10 mM dNTP, 1 µl 25 µM Truseq-i5-connector, 1 µl 25 µM Truseq-i7-connector (Supplementary Table 2), 0.6 µl 25 mM MgCl_2_, 0.3 µl KAPA enzyme (Roche), and 13.2 µl ddH_2_O. 1st round PCR was performed as: 72°C 5 min; 95°C 2 min; 13 cycles of 98°C 20 s, 65°C 30 s, 72°C 1 min; 72°C 5 min. 0.5 µl ExoI (NEB) was added into each well and the reaction was incubated at 37°C 1 hour followed by 72°C 20 min to deactivate ExoI. 2nd round PCR mix was prepared as follows: 2 µl 5xHiFi buffer, 0.3 µl 10 mM dNTP, 1 µl 25 µM Truseq P5, 1 µl 25 µM Truseq P7 (Supplementary Table 2), 0.2 µl 25 mM MgCl_2_, 0.15 µl KAPA enzyme, and 5.35 µl ddH_2_O. 2nd round PCR was performed as: 95°C 2 min; 5 cycles of 98°C 20 s, 60°C 30 s, 72°C 1 min; 72°C 5 min. The PCR product was purified with 0.9× AMPure XP beads (Beckman) followed by 0.45× + 0.45× double size selection. The final library product was eluted in 20 µl ddH_2_O and its concentration was measured with Qubit. Libraries may be sequenced with the standard recipe on the Nova-seq platform from Illumina or T7 and MGI 2000 platform.

### uCoTargetX

#### Reverse transcription

Rehydrated cells were washed twice with 0.01% TX-100 in 0.1%BSA-PBS for permeabilization. Cells were resuspended with 50 µl reverse transcription mix, which contains 1.25 µl 1 M Tris-HCl pH=8.3, 0.3 µl 5 M NaCl, 2.5 µl 10 mM GTP, 0.125 µl 1 M MgCl_2_, 4 µl 0.1 M DTT, 2.5 µl 10 mM dNTPs, 2.5 µl 200 U/μl Maxima H minus, 0.5 µl Takara RNase inhibitor, 1 µl 100 μM TSO µl, and 30.625 µl ddH_2_O. Incubate the system at 50°C for 10 minutes then 3 cycles of 8°C for 12 s, 15°C for 45 s, 20°C for 45 s, 30°C for 30 s, 42°C for 2 min and 50°C for 3 min followed by a final step at 50°C for 5 min. After reverse transcription, wash cells twice with NSB and resuspend cells with 50 μl NSB. RNA from RNA/cDNA hybrids were removed by 50 μg/ml RNase A (thermo) digestion at 37°C for 30 min.

#### Antibody targeted tagmentation and 2-round ligation

Cells were incubated with Antibody-PAT-T5&T7 complex in 100 μl Wash-buffer-TX-high salt plus 2 mM EDTA at room temperature for 1 hour. After incubation, cells were washed 3 times with Wash-buffer-TX-high salt to remove free unbound Antibody-PAT-T7 complex and resuspended with 50 μl Reaction buffer. The reaction was incubated at 37°C for 1 hour at a thermal cycler. Hot lid is set at 40°C. The reaction was stopped by incubating cells in 180 µl Wash-buffer-TX-high salt with 5 mM EDTA at room temperature for 5 min. The above steps were repeated N times for the other Antibody-PAT-T5&T7 complex. After targeted tagmentation, cells were then submitted for 2-round ligation procedures just like uCoTarget. 3,000-5,000 cells were aliquoted to 96-well plates as many as possible for lysis. Lysis and Proteinase K deactivation protocol was as above.

#### Pre-amplification

Pre-amplification mix was prepared as follows: 5 µl cell lysate, 10 µl 5xHiFi buffer, 1 µl 10 mM dNTP, 2 µl 10 µM Biotin-IS primer, 4 µl 10 μM Truseq-i5-connector, 2 µl 10 μM Truseq-i7-connector, 1 µl 25 mM MgCl_2_, 0.5 µl KAPA enzyme, and 24.5 µl. Pre-amplification was performed as: 72°C 5 min; 95°C 2 min; 6 cycles of 98°C 20 s, 65°C 30 s, 72°C 2 min 30 s; 72°C 5 min. Pre-amplification product was purified with 0.9× AMPure XP beads and eluted in 25 µl ddH_2_O. Activated C1 beads was prepared as follows: 1 µl original C1 beads was washed twice with 2xB&W buffer (10 mM Tris-HCl pH 7.5, 2 M NaCl, 1 mM EDTA and 0.05% Tween 20) and resuspended with 25 µl 2xB&W buffer. The 25 µl activated C1 beads was added to 25 µl pre-amplified DNA/RNA products and incubate at room temperature for 30 min with rotation. C1 beads (RNA part) and supernatant (DNA part) were divided by a magnetic stand.

#### DNA-part library preparation

DNA part was purified with 0.45× + 0.45× AMPure beads for size selection and eluted in 10 µl ddH_2_O. Indexed PCR mix was prepared as follows: 10 µl 5xHiFi buffer, 1 µl 10 mM dNTP, 1 µl 25 µM Truseq P5, 1 µl 25 µM Truseq P7, 1 µl 25 mM MgCl_2_, 0.5 µl KAPA enzyme, 35.5 µl ddH_2_O. PCR was performed as: 95°C 2 min; 9 cycles of 98°C 20s, 65°C 30s, 72°C 1 min; 72°C 5 min. The PCR product was purified with 0.9× AMPure XP beads followed by 0.45× + 0.45× double size selection. The final library product was eluted in 20 µl ddH_2_O and its concentration was measured with Qubit. Libraries may be sequenced with the standard recipe on the Nova-seq platform from Illumina or T7 and MGI 2000 platform.

#### RNA-part library preparation

C1 beads was briefly washed with 180 µl Wash Buffer (+0.05% TX-100) twice. 50 µl re-amplification PCR mix was mixed with C1 beads, containing 10 µl 5xHiFi buffer, 1 µl 10 mM dNTP, 2.5 µl 10 µM IS primer (no biotin), 2.5 µl 10 µM Truseq-i5-connector, 1 µl 25 mM MgCl_2_, 0.5 µl KAPA enzyme, 32.5 µl ddH_2_O. Re-amplification was performed as: 95°C 2 min; 9 cycles of 98°C 20s, 65°C 30s, 72°C 2 min 30 s; 72°C 5 min. Re-amplified cDNA was purified with 0.9×AMPure XP beads and eluted with 16 µl ddH_2_O. The concentration of cDNA was quantified by Qubit. 50 ng cDNA was tagmented by Tn5-MEB at 55°C 5 min using the following system: 50 ng cDNA, 1 µl 500 nM Tn5-MEB, 1.5 µl 5×TAPS-MgCl_2_ (50 mM TAPS-NaOH, pH=8.3, 25 mM MgCl_2_), ddH_2_O to 7.5 µl. Indexed PCR mix was prepared as follows: 7.5 µl tagmented cDNA, 10 µl 5xHiFi buffer, 1 µl 10 mM dNTP, 1 µl 25 µM Truseq P5, 1 µl 25 µM Nextera i7, 0.5 µl KAPA enzyme, 29 µl ddH_2_O. Indexed PCR was performed as: 72°C 5 min; 95°C 2 min; 7 cycles of 98°C 20s, 65°C 30s, 72°C 1 min; 72°C 5 min. The PCR product was purified with 0.8× AMPure XP beads. The final library product was eluted in 20 µl ddH_2_O and its concentration was measured with Qubit. Libraries may be sequenced with the standard recipe on the Nova-seq platform from Illumina or T7 and MGI 2000 platform.

#### CoTarget data processing

CoTarget data were processed to generate unique and non-duplicated reads as previously described^2^. Briefly, we evaluated the quality of CoTarget sequencing data by FastQC (version 0.11.5). Low-quality bases and adapters were removed by cutadapt (version 1.11) with following parameters: -q 20 -O 10–trim-n -m 30– max-n 0.1. Clean paired-end CoTarget reads were then mapped to human and mouse reference genome hg19 and mm10 using Bowtie2 (version 2.2.9)^6^. The mapped reads with MAPQ greater than 30 were considered as uniquely mapped reads, which were sorted using Samtools (version 1.9) and used for subsequent analyses. PCR duplicates were removed by Picard (version 2.2.4) (http://broadinstitute.github.io/picard) with default parameters. Only uniquely mapped, non-duplicated reads were used for peak calling by MACS2 (version 2.1.1)^7^.

#### Receiver Operating Characteristic (ROC) curve

We performed ROC analysis to evaluate the CoTarget data quality. We chose K562 H3K27ac and H3K27me3 ChIP-seq peaks as gold standard from ENCODE database^8^. The peaks within regions of ± 5 kb of gene TSS were defined as standard positive peaks. True positive peaks were defined as CoTarget peaks overlapping with standard positive peaks. We next defined regions in TSS ± 5 kb which have no overlap with bulk ChIP-seq peak as standard false positive regions. False positive peaks were defined as CoTarget peaks not overlapping with standard positive peaks. We calculated true positive rate (TPR) as the number of true positive peaks divided by the number of standard positive peaks and false positive rate (FPR) as the number of false positive peaks divided by the number of standard false positive regions. To plot the ROC curves, we set a series of p value cut-off for peaks and generated a vector comprising TPR and FPR. For each group with different cell numbers and methods, we plotted ROC curve and calculated Area Under Curve (AUC) in R (version 4.2.2).

#### Visualization and correlation analysis of uCoTarget data

We used bamCoverage function in deepTools (version 2.2.3)^9^ to calculate genome coverage in BAM files and generated track files (bigwig format) for uCoTarget data. For visualization of the signals of histone modifications at specific loci, bigwig files were uploaded into Integrative Genomics Viewer (version 2.3)^10^ together with corresponding reference tracks. To evaluate correlations of H3K27ac and H3K27me3 between CoTarget, *in situ* ChIP, and bulk ChIP-seq in K562, we calculated the normalized average scores in genome 5-kb bins. The correlation was calculated between different groups and plotted by deepTools (version 2.2.3).

#### Collision rate

To evaluate the efficiency of split-and-pool strategy for single-cell labeling of uCoTarget, we calculated the collision rate (i.e. the number of human and mouse collisions over the total number of cells) by performing H3K27ac-H3K27me3 uCoTarget experiments using the 1:1 mixture of human (K562) and mouse (V6.5) single cells. We generated scatter plots using the proportion of reads mapping to human or mouse genome in each barcode combination in Figure 2. Barcodes with <90% of aligned reads mapped to one species were classified as collisions^11^.

#### Single-cell uCoTarget data processing

The uCoTarget data were demultiplexed by custom scripts. Briefly, the analysis pipeline of uCoTarget data processing consisted of following steps: (i) Creating the single cell whitelist using UMI-tools (version 1.1.2)^12^ based on the molecular design of uCoTarget (Supplementary Figure 1) with the parameter “--set-cell-number=n”, where n is the true cells used in the uCoTarget experiments; (ii) Extracting the paired-end reads based on the single-cell whitelist; (iii) Mapping to the human or mouse reference genomes by bowtie2 (version 2.2.9)^6^; (iv) Keeping uniquely mapped reads and removing PCR duplicates as the performance in CoTarget data processing; (v) Adding cell barcode information to the bam files and generating single-cell bam files^13^.

#### Single-cell uCoTarget data processing

uCoTargetX DNA-part data processing was performed as described in the “uCoTarget data processing” section mentioned above. The uCoTargetX RNA-part data were demultiplexed by custom scripts. Briefly, (i) Creating the single cell whitelist using UMI-tools (version 1.1.2)^12^ based on the molecular design of uCoTargetX with the parameter “--set-cell-number=n”, where n is the true cells used in the uCoTarget experiments; (ii) Extracting the paired-end reads based on the single-cell whitelist; (iii) Mapping to the human (hg19) or mouse (mm10) reference genomes using Hisat2 (version 2.0.4); (iv) The gene expression level was quantified by UMI-tools (version 1.1.2).

#### Single-cell clustering analysis

Following Single-cell uCoTarget data processing above, we generated a peak-by-cell matrix for uCoTarget datasets using the “createcisTopicObjectFromBAM” function in cisTopic (version 0.3.0)^14^. Using the peak-by-cell matrix as an input, we performed dimensionality reduction using latent semantic indexing (LSI) algorithm by running the “RunTFIDF” and“RunSVD” function in Signac (version 1.8.0)^15^. LSI dimensions that were highly correlated with read depth were identified using “DepthCor” function in Signac and were excluded in downstream analysis. We next ran Signac/Seurat’s ‘RunUMAP’ function on the LSI dimensions to compute the UMAP embedding followed by clustering analyses using Signac/Seurat’s ‘FindNeighbors’ and ‘FindClusters’ at varying resolutions. Clusters were annotated by H3K27ac signals near known marker genes.

#### Cicero co-binding analysis

We calculated co-binding scores of single cells of different cell types for H3K4me3, H3K27ac, and H3K27me3 histone marks in our uCoTarget data using the R package Cicero (version 1.14.0)^16^ as previously described. Briefly, we first used the matrix from Seurat object and created cellDataSet. We next created a ‘cic-ero_cds’ using the “make_cicero_cds” function in Cicero based on corresponding UMAP coordinates. All peak-to-peak linkages were identified by running “run_cicero” with an input of a cicero_cds and genomic_coords. The linkages may be visualized around specific genomic regions.

#### ChromVAR TF motif score calculation

We calculated global TF activity score using R package chromVAR (version 1.18.0)^17^ with default parameters. Briefly, as input we used the raw counts for all peaks and the CISBP motif (from chromVAR motifs ‘human_pwms_v2’) matches within these peaks from motifmatchr. We then computed the GC bias-corrected deviation score using the “getBackgroundPeaks” and “computeDeviations” function in chromVAR. The deviation score was used to plot heatmap in R (version 4.2.2)

### Code availability

Codes for analysis of the figures are available at https://github.com/Helab-bioinformatics/uCoTarget.

### Data availability

Raw sequencing data have been deposited at the NCBI Gene Expression Omnibus (GEO) with the accession number GSE220193.

### Statistics and reproducibility

All uCoTarget experiments were independently performed at least twice. Other statistical analyses in this study were mainly performed between groups of peaks or segments of the genome. Statistical tests are indicated as in the figure legends.

## Notes

### Competing Interest Statement

The authors have declared no competing interest.

