## Supplementary Figures for "Massively parallel single-cell profiling of transcriptome and multiple epigenetic proteins in cell fate regulation"

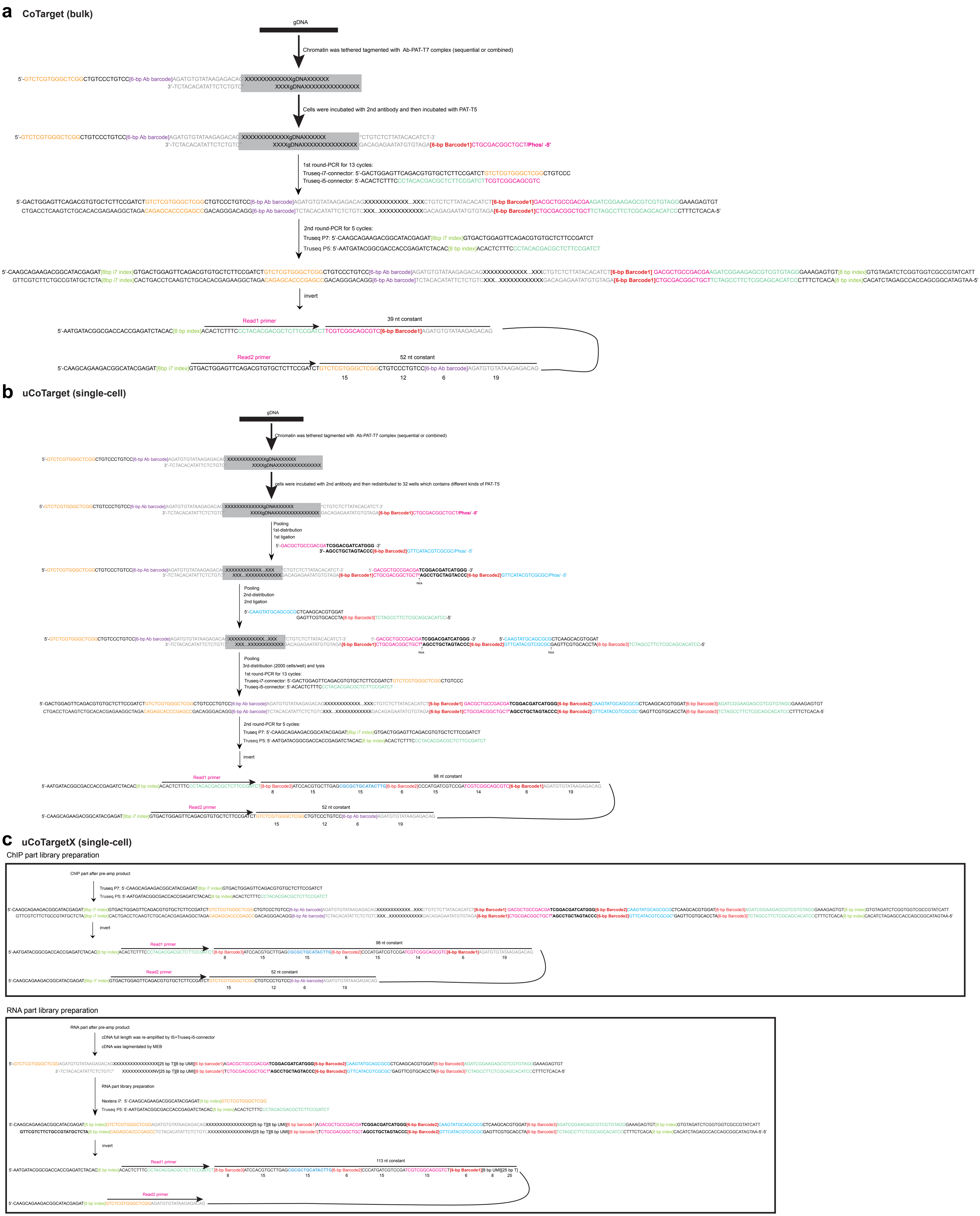


**Supplementary Figure 1. Overview of the molecular design of the library preparation of CoTarget, uCoTarget, and uCoTargetX.** (a-c) The molecular design of CoTarget (a) for bulk samples, uCoTarget (b), uCoTargetX (c) for single cells.

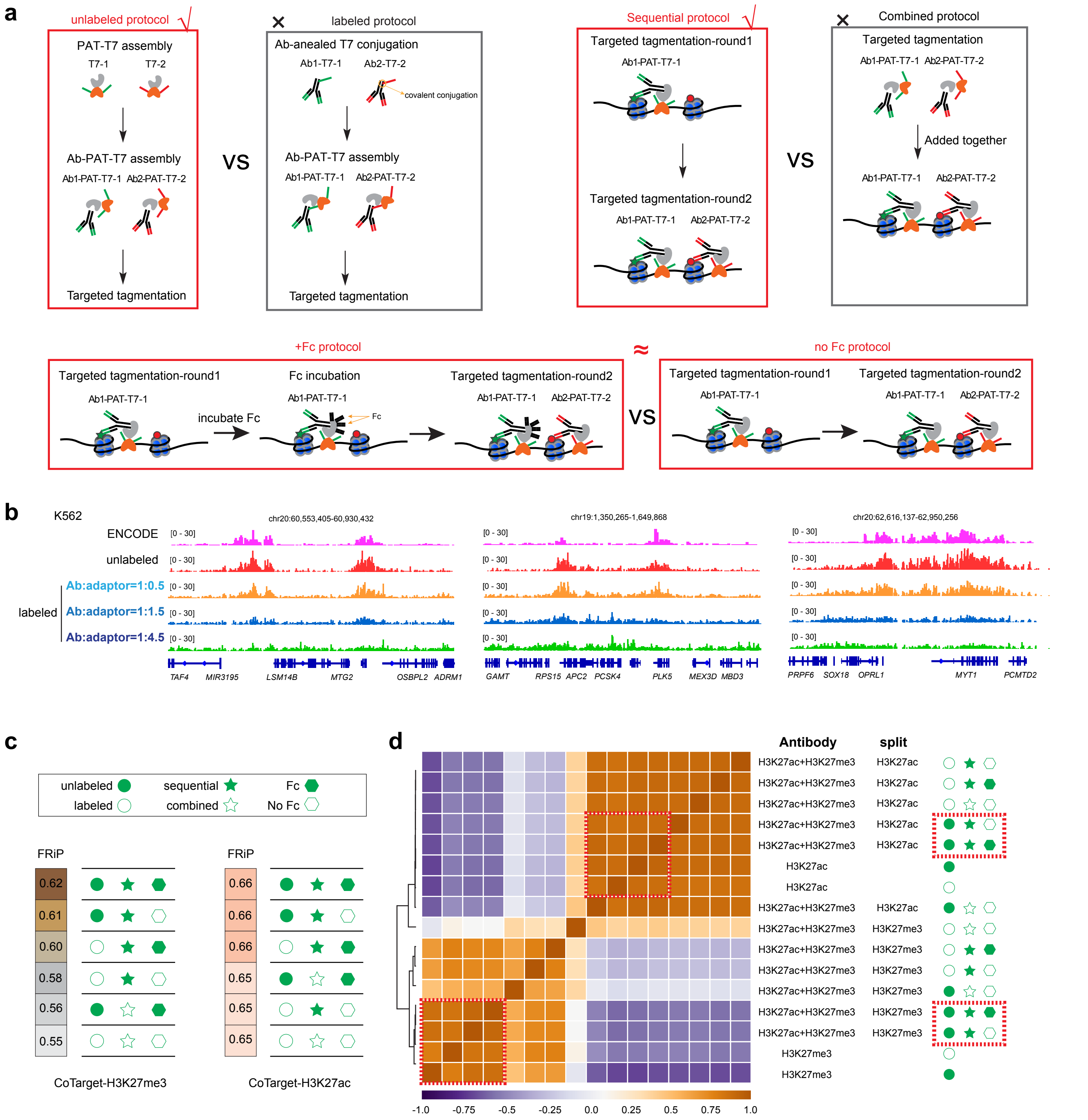


**Supplementary Figure 2. Optimization and benchmarking of key steps of the CoTarget procedure.**
(a) Optimization of the main steps of the CoTarget procedure. The conditions labeled with red box were selected in subsequent CoTarget expeperiments.
(b) Track view showing H3K27me3 CoTarget signals in K562 with different labeling protocol. For labeled groups, antibodies were first covalently conjugated with T7 adaptors with different mole ratio (1:0.5-1:4.5). Labeled antibodies were then incubated with PAT to form antibody-PAT-T7 complex for CoTarget experiments.
(c) Fraction of reads in peaks (FRiP) for different groups with indicated combination of experimental conditions.
(d) Heatmap showing correlation between different conditions. The experimental conditions that most correlated with corresponding in situ bulk ChIP data were labeled with dash red boxes.

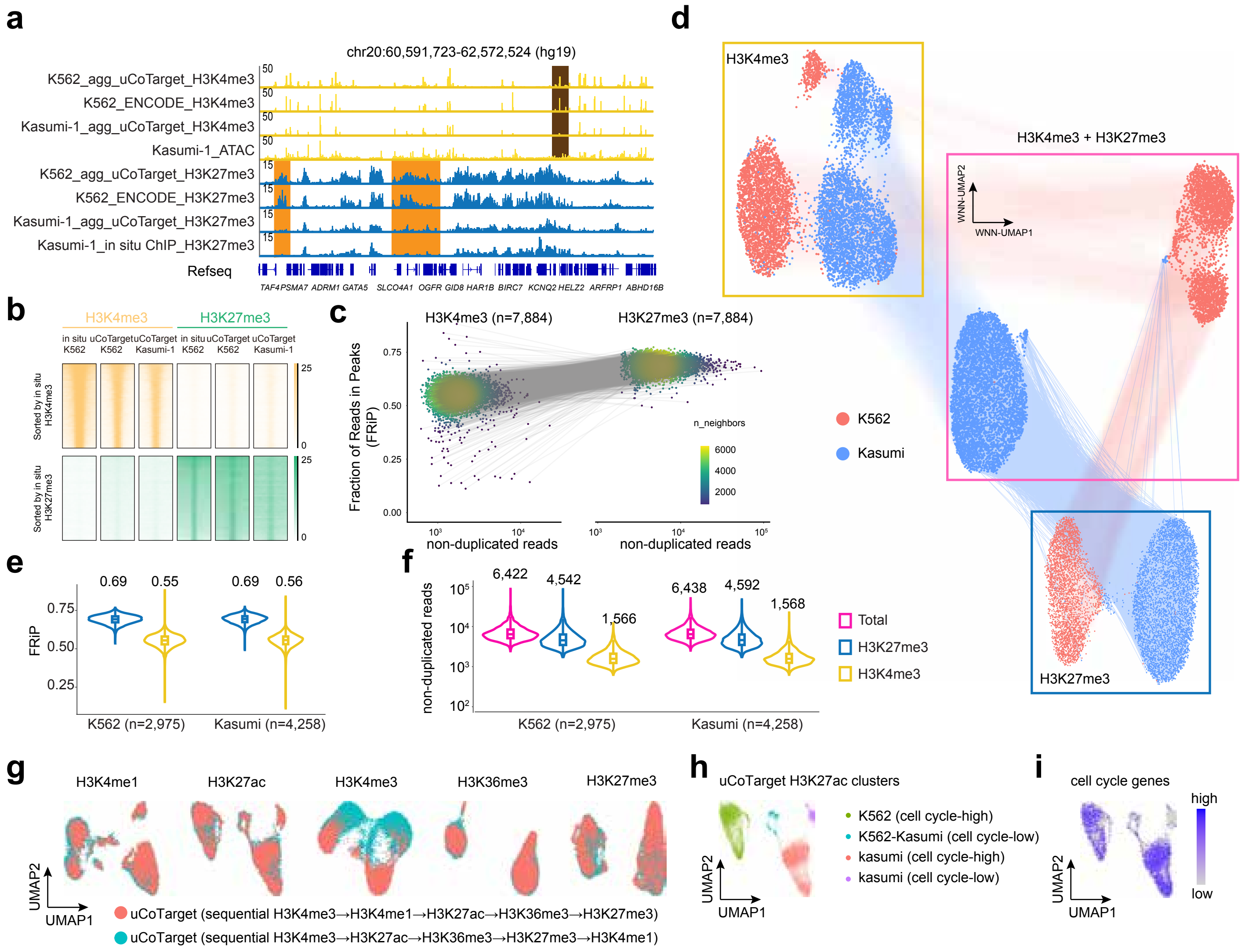


**Supplementary Figure 3. Quality control of uCoTarget data in K562 and Kasumi-1 cells.**

(a) The aggregate track view of uCoTarget profiling of two histone modifications.

(b) Heatmap showing H3K4me3 (up) and H3K27me3 (bottom) signals in uCoTarget aggregate data and in situ ChIP data at peak regions.

(c) Scatter plots with cell-cell linkage showing the non-duplicated reads and the fraction of reads in peaks (FRiP)of each cell. Color bar represents the cell density.

(d) Connected UMAP visualization of single modality and WNN integration of 2 modalities. Dots connected by lines represent the same cells between different visulazations.

(e) Violin plots showing the FRiP of each single cells (blue: H3K27me3, yellow: H3K4me3). The median FRiP value was displayed on the top of each violin.

(f) Violin plots depicting the distribution of non-duplicated reads per cells of each modalities (blue: H3K27me3, yellow: H3K4me3) and total reads detected in a cell (purple). The median read number was shown on the top.

(g) UMAP visualization of single cells from using different modification information. Color represents the sequential order of five histone modifications co-profiled.

(h-i) UMAP visualization of cell heterogenities using uCoTarget H3K27ac clusters (h) and the distribution of the H3K27ac signals on cell cycle signature genes (i).

**a**

| Methods | Specific fusion proteins | Special preparation | Throughput | Specialized device | # target | TFs | Transcriptome |
|---|---|---|---|---|---|---|---|
| multi-CUT&Tag (Gopalan et al., 2021, Mol. Cell) | No | His-tag purification for Ab·Tn5 complex | High | 10X ATAC platform | 2 | No | No |
| NTT-seq (Stuart et al., 2022, Nat Biotechnol.) | Species specific secondary nanobody-Tn5 | No | High | 10X ATAC platform | 3 | No | No |
| Nano-CT (Bartosovic et al., 2022, Nat Biotechnol.) | Species specific secondary nanobody-Tn5 | No | High | 10X ATAC platform | 3 | No | No |
| MulTI-Tag (Meers et al., 2022, Nat Biotechnol.) | No | Ab·adaptor covalent conjugation | High | ICELL8+microwells | 3 | No | No |
| **uCoTargetX (This study)** | No | No | **Ultra high** | No | **5** | √ | √ |

**b**

| Methods | Sample | Histone marks or TFs | # single cells | Non-duplicated reads |
|---|---|---|---|---|
| multi-CUT&Tag (Gopalan et al., 2021, Mol. Cell) | Mixture of mESCs and mTSCs | H3K27ac+H3K27me3 | 1,949 | not reported (200 unique reads as cut off for quality control) |
| NTT-seq (Stuart et al., 2022, Nat Biotechnol.) | Mixture of K562 and HEK293 | H3K27me3+H3K27ac+RNAPII | 8,617 | 743 (H3K27me3), 382 (H3K27ac), 542 (RNAPII) |
| | PBMCs | H3K27me3+H3K27ac+protein | 4,684 | 2,854 (H3K27me3), 412 (H3K27ac), 690 (ADT counts) |
| | BMMCs | H3K27me3+H3K27ac | 5,236 | 1,217 (H3K27me3), 326 (H3K27ac) |
| Nano-CT (Bartosovic et al., 2022, Nat Biotechnol.) | P19 mouse brain | H3K27me3+H3K27ac | 11,981 | ~3,000 (H3K27me3), 10,000 (H3K27ac) |
| | | H3K27me3+H3K27ac+ATAC | 4,434 | ~3,000 (H3K27me3), 10,000 (H3K27ac), 3,000 (ATAC) |
| MulTI-Tag (Meers et al., 2022, Nat Biotechnol.) | K562, H1, and a mixture of them | H3K27me3+H3K36me3 | 21,548 | 1,416 (H3K27me3), 436 (H3K36me3) |
| | H1 | H3K27me3+H3K4me2+ H3K36me3 | 373 | 2,765 (H3K27me3), 287 (H3K4me3), 358 (H3K36me3) |
| | K562 | | 372 | 3,568 (H3K27me3), 193 (H3K4me3), 217 (H3K36me3) |
| | Differentiated H1 | H3K27me3+H3K36me3+ H3K4me1 | 387 (H1)<br>827 (endoderm)<br>737 (mesoderm)<br>337 (ectoderm) | 1,716 (H3K27me3), 611 (H3K36me3), 144 (H3K4me1)<br>1,534 (H3K27me3), 381 (H3K36me3), 170 (H3K4me1)<br>2,712 (H3K27me3), 519 (H3K36me3), 153 (H3K4me1)<br>415 (H3K27me3), 423 (H3K36me3), 170 (H3K4me1) |
| **uCoTargetX (This study)** | K562 and Kasumi | H3K27me3+H3K4me3 | 7,884 | 4,489 (H3K27me3), 1,535 (H3K4me3) |
| | | H3K4me3+H3K4me1+H3K27ac+ H3K36me3+H3K27me3 | 19,860 | 3,030 (H3K4me1), 2,484 (H3K27me3), 1,937 (H3K27ac), 1,839 (H3K4me3), 1,377 (H3K36me3) |
| | Differentiation from hESC to HSPC | H3K27me3+H3K27ac+H3K4me3 | 15,731 | 11,437 (H3K27ac), 6,078 (H3K27me3), 1,851 (H3K4me3) |
| | | H3K27ac+RUNX1+H3K27me3 | 10,687 | 6,894 (H3K27ac), 2,557 (H3K27me3), 1,390 (RUNX1) |

**c**

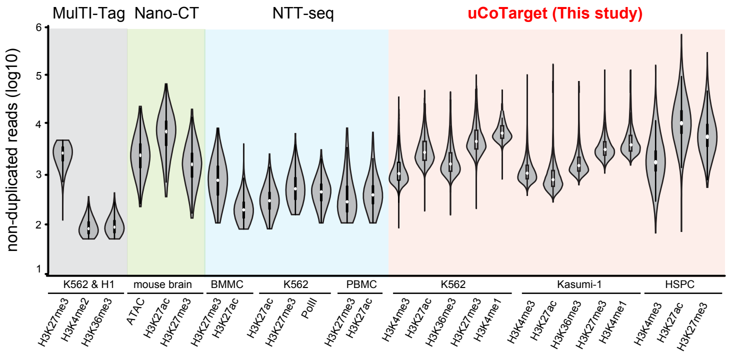


**d**

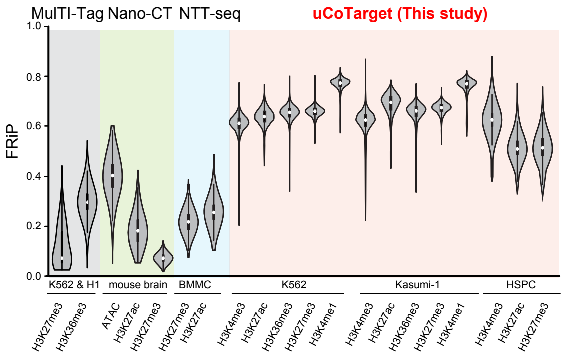


**Supplementary Figure 4. Comparison of uCoTarget with exsiting methods.**
(a) Comparison of experimental strategy and applications.
(b) Data quality assessment of uCoTarget and other methods on different samples.
(c-d) Violin plot showing the distribution of non-duplicated reads (c) and fraction of reads (FRIP) (d) in different single-cell methods, including MulTI-Tag, Nano-CT, NTT-seq, and uCoTarget. Public data were downloaded from GSE179756, GSE198467, and GSE212588 for MulTI-Tag, Nano-CT, and NTT-seq respectively. The boxes in violin plots here indicate upper and lower quartiles (25th and 75th percentiles). Since cell-sepecific barcode sequences of multi-CUT&Tag datasets were trimmed off in raw data as provided in GEO database, multi-CUT&Tag datasets were not analyzed here.

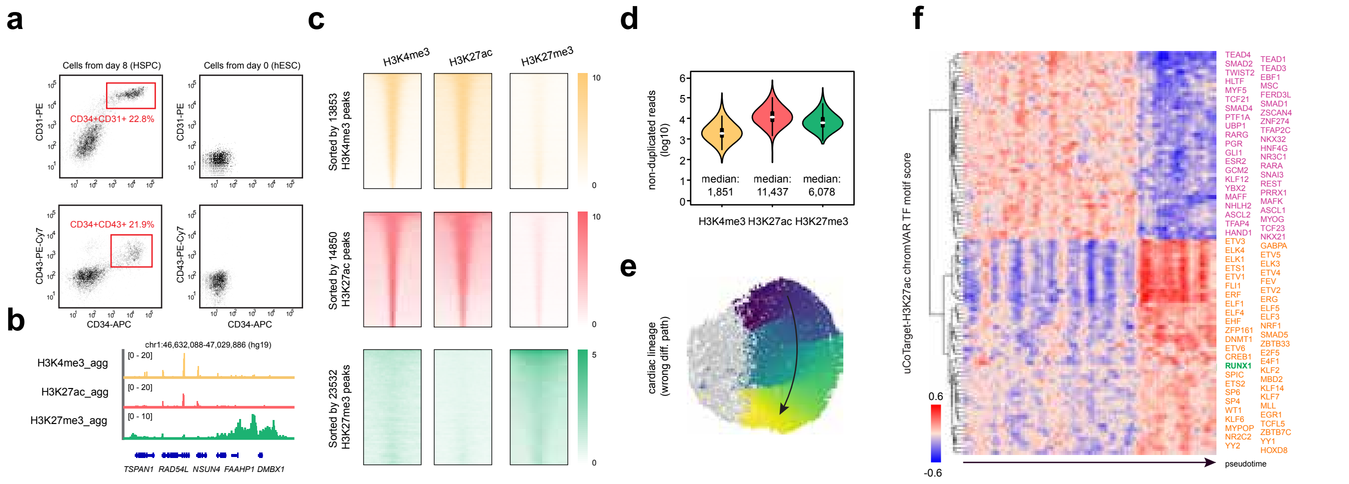


**Supplementary Figure 5. uCoTarget profiling of H3K27me3, H3K27ac, and H3K4me3 during HSPCs differentiation.**

(a) Flow cytometry plots show expression of CD34 (marker gene for HSPC), CD31 (marker gene for endothelial cell), and CD43 (marker gene for pre-HSC) in differentiation day 8 samples and undifferentiated day 0 samples.

(b) Track view showing aggregate H3K4me3, H3K27ac and H3K27me3 signals at representive loci.

(c) Heatmap showing H3K4me3 (top), H3K27ac (middle) and H3K27me3 (bottom) signals in different modalities of uCoTarget data.

(d) Violin plots showing non-duplicated reads per cell for each histone modification. The boxes in violin plots indicate upper and lower quartiles (25th and 75th percentiles).

(e) Pseudotime analysis of cardiac lineages. The color from dark blue to yellow indicates pseudotime from early to late. For cardiac lineages, the hematopoietic cells colored by grey were not used for pseudotime analysis here.

(f) ChromVAR identifying TF dynamics along HSPC differentiation path.

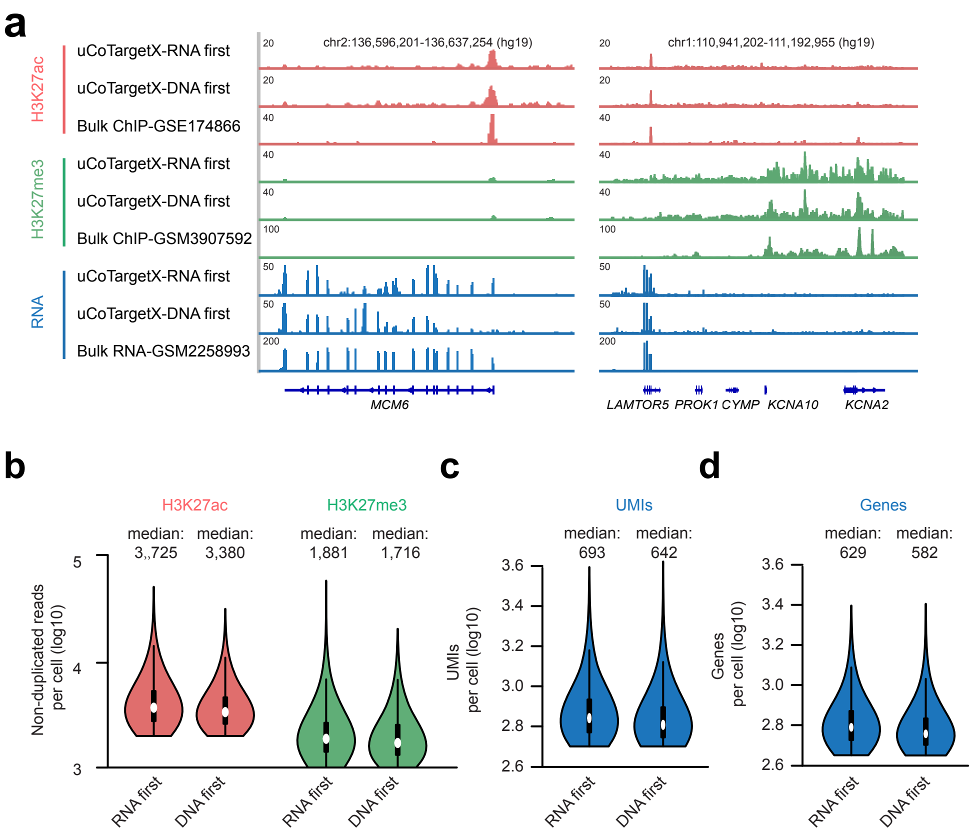


**Supplementary Figure 6. Optimization and benchmarking of key steps of the CoTargetX procedure in 293AD cells.**

(a) Track view displaying both RNA, H3K27ac and H3K27me3 signals on representative gene loci in 293AD cells for the condition of dealing with DNA or RNA first in the uCoTargetX protocol.

(b-d) Violin plots showing non-duplicated reads per cell (b) detected UMI (c) and  genes (d) per cell in 293AD cells for the condition of dealing with DNA or RNA first in the uCoTargetX protocol. The boxes in violin plots indicate upper and lower quartiles (25th and 75th percentiles).
